# Late treatment initiation leads to reduced antiviral potency

**DOI:** 10.1101/2025.09.24.678199

**Authors:** Xuanlin Liu, Evelyn J. Franco, Sean N. Avedissian, Kaley C. Hanrahan, Jeremie Guedj, J. G. C. van Hasselt, Ashley N. Brown, Anne-Grete Märtson

## Abstract

The timing of initiation is critical in antiviral treatment and viral dynamic (VD) modeling is a powerful tool to study the within-host viral load changes and evaluate antiviral treatment effects using mathematical equations. Previous simulation studies have shown that early treatment initiation is critical to maximize the therapeutic response in antiviral treatment in an acute viral infection such as influenza and SARS-CoV-2. A recent experimental study demonstrated that late therapy initiation can lead to diminished antiviral potency. However, most VD model simulations with varying treatment initiation time accounted only for the effect of initiation condition (i.e., state of different cell populations when the therapy started), the loss of drug potency has been under-investigated. This may overestimate the antiviral effect, potentially resulting in suboptimal dose selection. To this end, we aimed to characterize relationship between the drug potency (EC_50_) and the timing of drug addition, using nirmatrelvir and GS-441524 against SARS-CoV-2 as an example. Viral load data were obtained from *in vitro* experiments with various drug concentrations and treatment initiated between 0 to 3 days post infection. EC_50_ values were fitted for each treatment initiation group and were found to vary with the timing of treatment initiation in both drugs. Also, a VD model with time-varying EC_50_ provided better fits than a constant EC_50_ model (BIC = 1667.90 vs. 1677.84). Further simulations also indicated that a constant EC_50_ model overestimated the antiviral efficacy when treatment started late. These findings highlighted the importance of considering EC_50_ shift when optimizing dosage regimens for patients presenting late.

## Introduction

Severe acute respiratory coronavirus 2 (SARS-CoV-2) still causes severe infections in high risk patient groups, such as individuals with advanced age (≥65 years) and immunocompromised patients (1). Several drugs have been approved to treat SARS-CoV-2 infection: nirmatrelvir/ritonavir, remdesivir, and molnupiravir (2). Early treatment initiation is critical as late therapy initiation can result in reduced drug efficacy (3). Elucidating the relationship between drug efficacy and the time of therapy initiation enables us to optimize the dosing regimen in this scenario.

Several mathematical viral dynamic models have been developed using viral load data collected from either *in vitro* experiments or untreated infected patients (4–9). Many studies have tested the impact of timing of therapy initiation and dosage regimen on antiviral effectiveness based on model simulation (10–18). It has been established that early intervention is of vital importance to achieve ideal therapeutic effect (12, 14, 17, 18). Most of the published studies (10–18) predicted the viral load trajectories by combining a traditional target cell-limited (TCL) model with an antiviral therapy, which could reduce viral infection rate or production rate.

A classical TCL model assumed a constant half-maximal effective concentration (EC_50_) regardless of the progression of the viral infection when the therapy started. Nevertheless, it has been shown that the antiviral activity of molnupiravir decreased significantly when the treatment started late (19). Thus, the antiviral effect depends not only on the drug concentration, but also on the time of treatment initiation. However, existing models did not take the loss of antiviral activity due to late treatment initiation into account, potentially overestimating the antiviral effect when the treatment begins late in the course of infection.

In this study, we aimed to investigate the impact of late treatment initiation on the antiviral effect of nirmatrelvir (without ritonavir) and the active metabolite of remdesivir (GS-441524) against *in vitro* SARS-CoV-2 infection. We set up an *in vitro* study to investigate the viral dynamics under different time of treatment initiation. We also intended to evaluate the antiviral effect of various dosing regimens and treatment initiation times by *in silico* model simulation.

## Results

### *In vitro* experimental results

The antiviral activity of nirmatrelvir and GS-441524 was assessed by monitoring the replication curve of infectious viral titer. The experiment was repeated in two independent replicates for each drug under the same condition. The viral load profiles from different drug concentration groups were stratified by treatment initiation time and depicted respectively. The nirmatrelvir group showed good consistency across two replicates (Fig. 1a). Although large variability can be seen in the GS-441524 data between two replicates, the overall trend and outcome was similar (Fig. 1b). The *in vitro* infection experiments showed the fastest viral load decline occurred when therapy was started on day 0. Antiviral effect was limited for all doses when therapy was initiated from day 1 onwards, indicating the drug antiviral activity was suppressed due to delayed therapy initiation, especially in high concentration groups.

**FIG 1:**
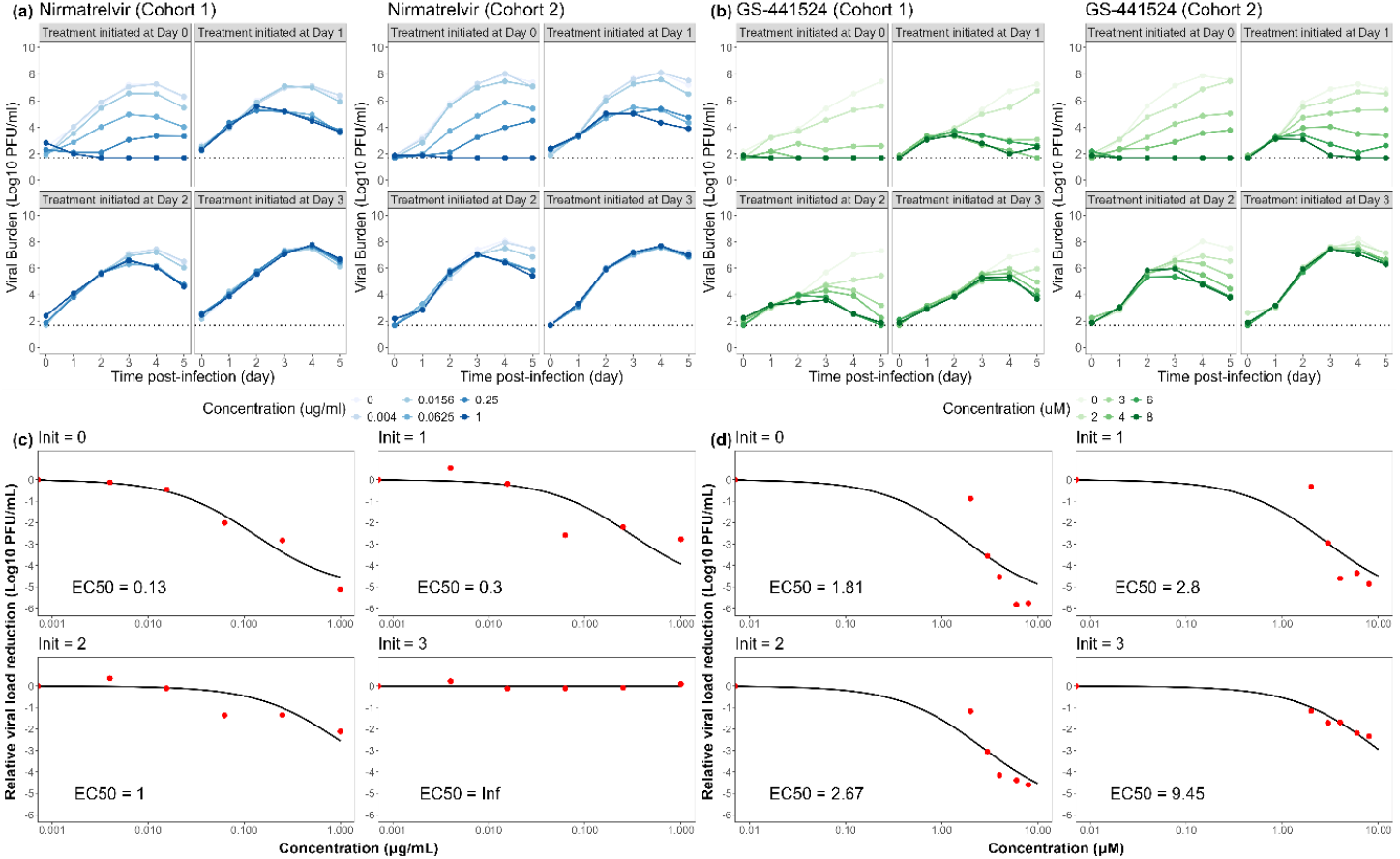
Replication curve of infectious viral load titer of (a) nirmatrelvir and (b) GS-441524 against *in vitro* SARS-CoV-2 infection in different concentration groups with varying treatment initiation times (i.e., 0-, 1-, 2-, 3-DPI). Each cohort means an independent experiment. Experiment-based EC_50_ of (c) nirmatrelvir and (d) GS-441524 at 5-DPI increased along with the delaying treatment initiation. The red points are observed data calculated from experimental results. The black line is the fitting curve using a sigmoidal E_max_ function. Black dotted thin lines represent the lower limit of quantification (100 PFU/mL). Abbreviations: EC_50_, apparent effective concentration at 50%; DPI, days post infection

The experiment-based EC_50_ of viral replication inhibition was derived from the viral load reduction relative to the control group. The fitting results showed that the EC_50_ increased when the treatment was further delayed. The fitted EC_50_ increased from 0.13 μg/mL to infinite for nirmatrelvir (Fig. 1c). As for GS-441524 (Fig. 1d), there was no significant difference in EC_50_ when the treatment started at 0-, 1-, 2-day post infection (DPI), which were 1.81 μM, 2.8 μM, and 2.67 μM respectively. However, the EC_50_ increased to 9.45 μM when the treatment started at 3-DPI.

### Viral dynamic modeling

Two target-cell limited (TCL) models with either single EC_50_ or multiple EC_50_ were developed and compared to see which model can better describe the experimental data. The viral dynamic parameters (i.e., viral infection rate *β* and viral production rate *ρ*) aligned well between the single EC_50_ model and the multiple EC_50_ model (Table 1)

**TABLE 1:** Parameter estimation of the TCL models with single and multiple EC_50_.

| Parameter (unit) | Single EC<sub>50</sub><br>Estimate (95% CI) | Multiple EC<sub>50</sub><br>Estimate (95% CI) |
| --- | --- | --- |
| β (1/PFU/day) | 3.73×10<sup>-8</sup> (2.1×10<sup>-8</sup>, 6.62×10<sup>-8</sup>) | 4.12×10<sup>-8</sup> (2.65×10<sup>-8</sup>, 6.41×10<sup>-8</sup>) |
| δ (1/day) | 2.58 fixed | 2.58 fixed |
| ρ (PFU/cell/day) | 1310 (726, 2370) | 1200 (763, 1900) |
| c (1/day) | 10 fixed | 10 fixed |
| E<sub>max</sub> | 1 fixed | 1 fixed |
| Nirmatrelvir EC<sub>50</sub> (µg/mL) | 0.26 (0.18, 0.37) | θ<sub>0</sub> = 0.17 (0.11, 0.26)<br>θ<sub>1</sub> = 0.48 (0.24, 0.96)<br>θ<sub>2</sub> = 0.64 (0.14, 2.92)<br>θ<sub>3</sub> = 16.9 (NA) |
| GS-441254 EC<sub>50</sub> (µM) | 1.95 (1.57, 2.41) | θ<sub>0</sub> = 2.32 (1.85, 2.91)<br>θ<sub>1</sub> = 1.92 (1.44, 2.57)<br>θ<sub>2</sub> = 1.00 (0.64, 1.59)<br>θ<sub>3</sub> = 1.59 (0.49, 5.14) |
| BIC | 1677.84 | 1667.90 |

Abbreviations: β, viral infection rate; δ, death rate of infected cells; ρ, viral production; c, viral clearance; E<sub>max</sub>, maximal antiviral effect; EC<sub>50</sub>, drug concentration required to reach 50% of the maximal antiviral effect; θ<sub>n</sub>, EC<sub>50</sub> when drug was added at n-day post infection; PFU, plaque forming unit; CI, confidence interval; BIC, Bayesian information criterion.

In the single EC_50_ model, the EC_50_ of nirmatrelvir and GS-441254 were 0.26 μg/mL and 1.95 μM respectively. On the other hand, a clear EC_50_ shift can be seen in the multiple EC_50_ model: the estimated nirmatrelvir EC_50_ values were 0.17, 0.48, 0.64 and 16.9 μg/mL when the drug added at 0-, 1-, 2-, 3-DPI respectively. The EC_50_ at 3-DPI cannot be estimated accurately as it has a wide range of 95% CI, which was similar to the experiment-based EC_50_ at 3-DPI. However, there was no clear trend in GS-441254 EC_50_ values.

The predictive performance of the single EC_50_ model and the multiple EC_50_ model were compared in Figure 2. In general, model with multiple EC_50_ (BIC = 1667.90) had a lower BIC than the model with single EC_50_ (BIC = 1677.84). Both models fit nirmatrelvir data (Fig. 2a and Fig. 2b) better than GS-441254 data (Fig. 2c and Fig. 2d), but the pattern of how the EC_50_ shift influencing the model was similar among all the study cohorts. In general, there was no significant difference between the two models in low concentration groups (nirmatrelvir concentration below 0.25 μg/mL or GS-441254 concentration below 6 μM). In nirmatrelvir high concentration groups (0.25 μg/mL and 1 μg/mL), the single EC_50_ model underestimated the viral load when the treatment started from 1-DPI onward, whereas the multiple EC_50_ model had a better fitting comparatively. Similar pattern can also be seen in GS-441254 high concentration group (6 μM and 8 μM), but the multiple EC_50_ model only had a better prediction when treatment started at 3-DPI.

**FIG 2:**
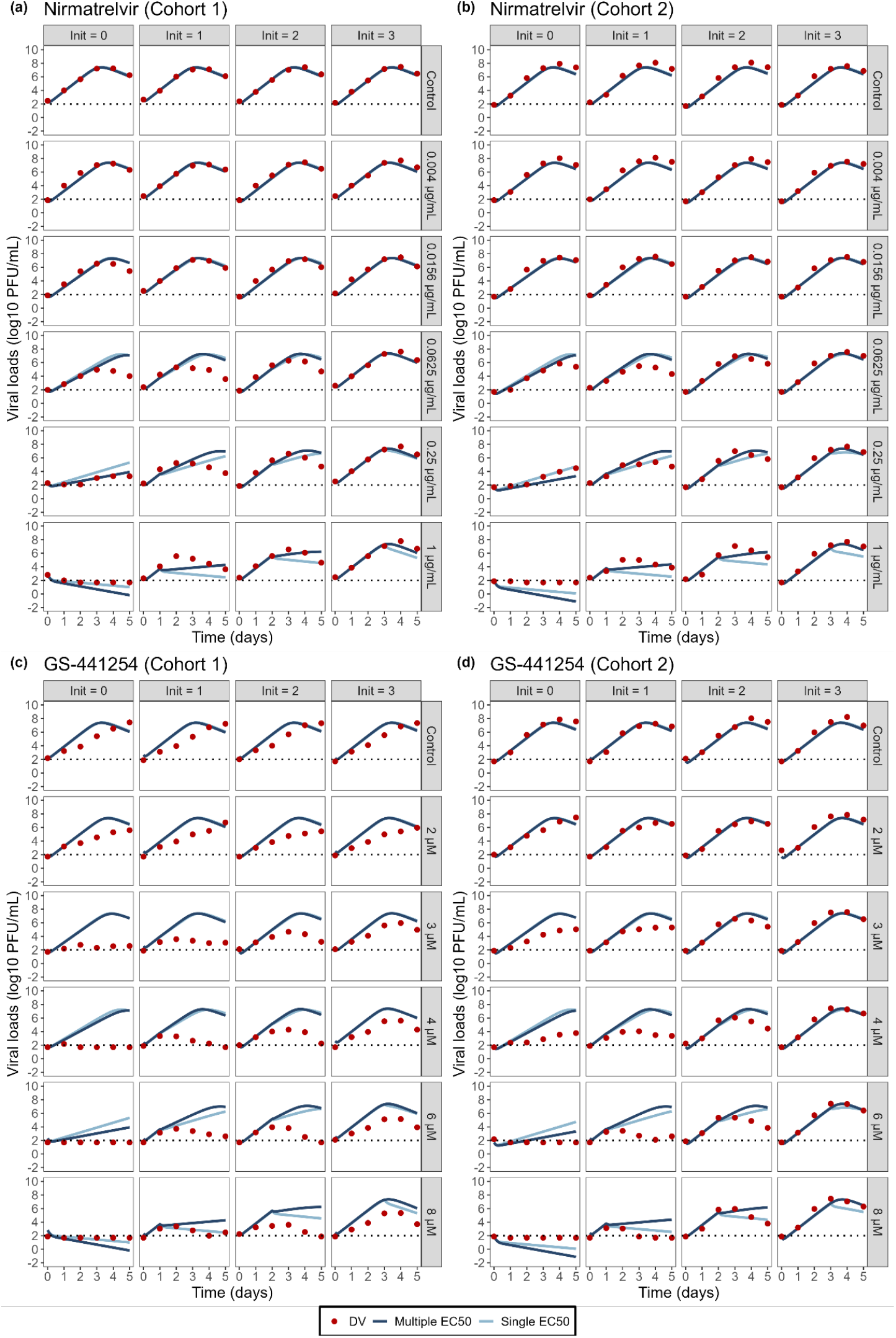
Individual prediction (IPRED) vs. observation (DV) of nirmatrelvir (a and b) and GS-441254 (c and d). The results are stratified on concentration and time of treatment initiation. The red solid points are observed viral load data from experiments. Dark blue solid lines represent the prediction from the model with multiple EC_50_. Light blue solid lines represent the prediction from the model with single EC_50_. Black dotted thin lines represent the lower limit of quantification (100 PFU/mL).

### Simulations of nirmatrelvir treatment in different scenarios

The standard nirmatrelvir treatment applied for *in vitro* simulation was 300 mg every 12 hours for 5 days. The plasma concentration time course of nirmatrelvir was generated using a published popPK model(18) (Fig. S1).

The influence of treatment initiation time was first tested. The dynamic of virus and different cells were simulated with varying time of treatment initiation: at 0-, 1-, 2-, 3-DPI, in comparison with the non-treatment group as control. Compared with TCL model with multiple EC_50_, the TCL model with single EC_50_ underestimated the viral load in when treatment started later than 1-DPI (Fig. 3a). However, the two models had no big differences when predicting the dynamics of target cells (Fig. 3b) and infected cells (Fig. 3c) when the treatment initiated no later than 2-DPI. While the TCL model with single EC_50_ had a higher prediction of target cell ratio and a lower prediction of infected cell ratio when the drug was given at 3-DPI, suggesting that the antiviral activity of nirmatrelvir was largely overestimated.

**FIG 3:**
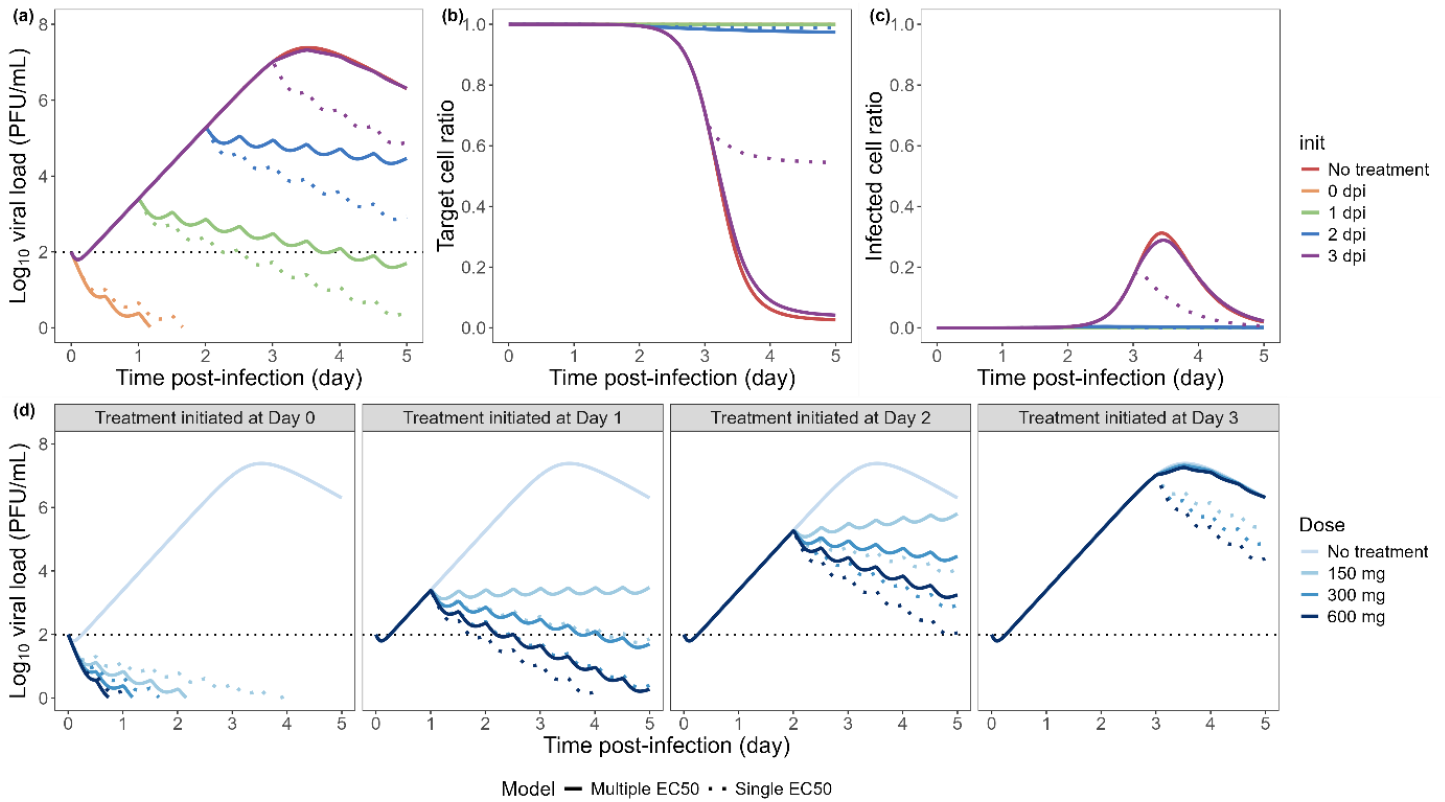
Simulated dynamics of virus (a), target cells (b) and infected cells (c) with standard nirmatrelvir treatment (300 mg q12h for 5 days) starting on 0-, 1-, 2-, 3-day post infection (DPI). (d) Simulated viral load trajectories with different dosing regimen (150mg, 300mg, 600 mg) when treatment starts at 0-, 1-, 2-, 3-DPI. Solid lines represent the prediction from the model with multiple EC_50_. Dotted lines represent the prediction from the model with single EC_50_. Black dotted thin lines represent the lower limit of quantification (100 PFU/mL).

On top of the treatment initiation time, the dose effect was also assessed by varying the standard dose within 2-fold. Similarly, single EC_50_ model exaggerated the antiviral activity of nirmatrelvir when treatment initiated after 1-DPI (Fig. 3d). When the treatments started at 0-DPI, all the dose groups were able to suppress the viral load below limit of quantification. When the treatments started at 1- and 2-DPI, the antiviral activity reduced in half dose group (150 mg) and standard dose group (300 mg), but it could be restored if we doubled the dose (600 mg). However, when the treatment was initiated at 3-DPI, there was no significant difference in the viral load trajectories among all concentration groups. Since all the target cells were infected at 3-DPI (Fig. 3b and Fig. 3c), it suggested that increasing dose would not provide extra benefit once the target cells were depleted.

## Discussion

In this study, we developed a hybrid method to investigate how the treatment initiation time influences the efficacy of nirmatrelvir and GS-441524, and verified the results using both experimental and modelling approaches.

For experimental approach, we conducted an *in vitro* antiviral experiment for nirmatrelvir (without ritonavir) and GS-441254 (the active metabolite of remdesivir) with static drug concentration. Each drug group had five concentration groups with a no-treatment control and the drug was added at 0-, 1-, 2-, 3-day post infection (DPI), corresponding to four treatment initiation times. We observed a reduced antiviral effectiveness when treatment started late (Fig. 4a). EC_50_ of nirmatrelvir and GS-441254 were then derived from the relative viral load reduction of each time of treatment initiation group. The experiment-based EC_50_ increased when the treatment started late.

**FIG 4:**
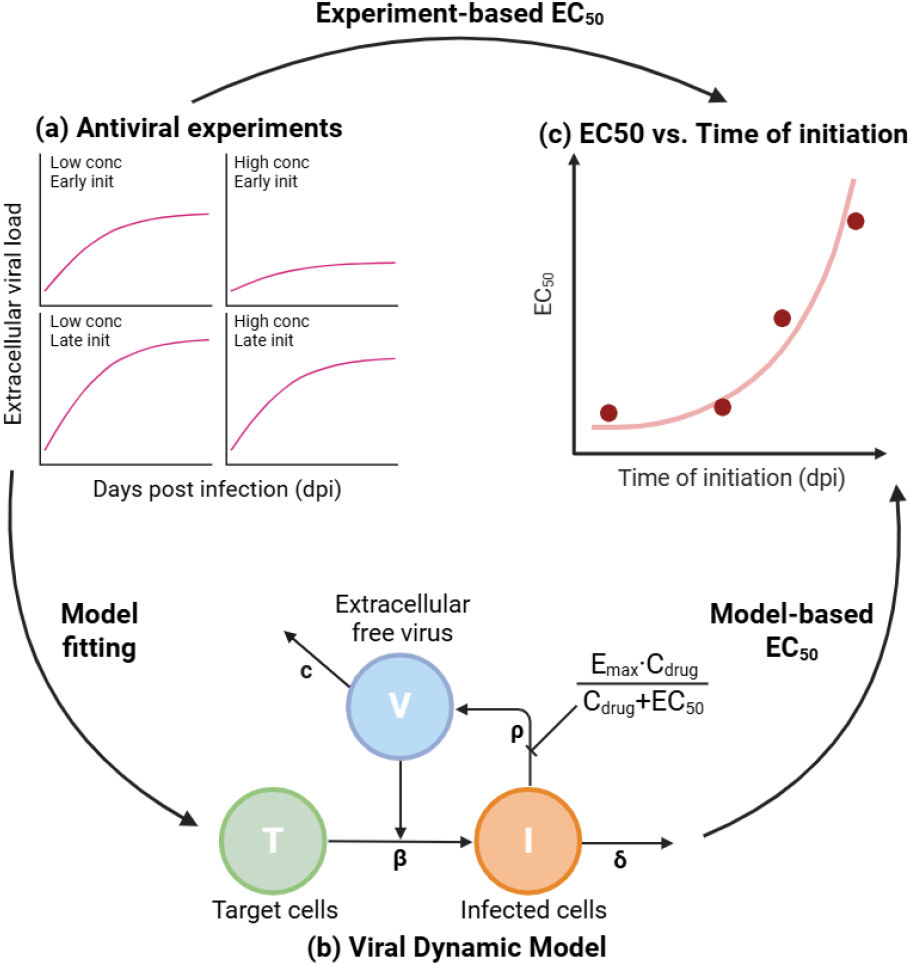
Diagram of the two-way hybrid verification of EC_50_ shift with late treatment initiation. (a) Antiviral experiments was performed for nirmatrelvir and GS-441524. Extracellular viral load from viral supernatant samples was measured daily for 5 days. Five static drug concentrations with four treatment initiation times were evaluated for each compound. EC_50_ was fitted separately for each treatment initiation time from the experimental results. (b) A target-cell limited (TCL) viral dynamic model incorporating a sigmoid E_max_ drug effect was developed. The EC_50_ of each treatment initiation time was estimated by fitting the viral load data to the TCL model. (c) The experiment - based EC_50_ and model-based EC_50_ were summarized to further investigate the relationship between EC_50_ and time of treatment initiation. Abbreviations: conc: drug concentration; init: time of treatment initiation; dpi: day post infection; δ, death rate of infected cells; ρ, release rate of intracellular virus; c, viral clearance; β, viral infection rate.

In terms of modelling approach, two TCL models with sigmoid E_max_ drug effect (Fig. 4b) were developed. One model used single EC_50_ shared by all the treatment initiation time groups, while the other one used multiple EC_50_ values Each time of treatment initiation group was assigned a specific EC_50_. The TCL model with multiple EC_50_ turned out to have a better predictive performance. The model-based EC_50_ of nirmatrelvir also increased when the treatment started late, while the EC_50_ shift was not clear for GS-441254. This could be resulted from the large variability between two replicates (Fig. 1b). Their baseline viral load trajectory was very different, the general viral load in cohort 1 is lower than cohort 2. The systematic differences could mask the potential EC_50_ shift during the modelling process and also lead to the suboptimal predictive performance when describing GS-441254 data.

By comparing the experiment-based EC_50_ and model-based EC_50_ with different time of initiation (Fig. 4c), we confirmed the EC_50_ was increasing along with the treatment initiation time.

To further assess the antiviral efficacy in dynamic drug concentration, both the single EC_50_ model and the multiple EC_50_ model were used to simulate viral load trajectories with nirmatrelvir treatment under different scenarios. When using the same dosage regimen but differing starting times of therapy, the simulation results suggested that there was a risk of overestimating drug antiviral activity when the treatment started late if a constant EC_50_ was applied in the model. However, this risk has long been under investigated.

Several SARS-CoV-2 viral dynamic models have been developed since the pandemic. Ke *et al*. identified the relationship between viral load and a person’s infectiousness in SARS-CoV-2(5). Zitzmann *et al*. investigated the key parameters affecting the robustness of data fitting in viral dynamic modelling (7). Recently, a more mechanistic model developed by Owens *et al*. was published, which was based on a larger dataset and considered the immune response (9). Gonçalves *et al*. (10), Kern *et al*. (12) and Zhang *et al*. (14) further evaluated different dosage regimens with varying treatment initiation times and suggested that early initiation time gives the most benefits to the patients. However, none of these models applied multiple or dynamic EC_50_ values.

Early studies have either focused on hypothetical antiviral treatment with an arbitrary drug efficacy (11, 13, 14) or repurposed drug use, most of which are no longer used clinically for SARS-CoV-2 treatment (12, 15, 17). Also, these predictions were not validated using clinical data from patients receiving delayed antiviral therapy. Thus, there is no evidence showing that a TCL model constant EC_50_ still applies when the treatment starts at late stage of infection.

We hereby confirmed that the EC_50_ would increase due to late treatment initiation, which consequently leads to loss of antiviral efficacy. Furthermore, we would like to propose that the loss of antiviral efficacy is not just a simple time-dependent effect, but rather the progressive accumulation of intracellular viral load. The importance of intracellular level of infection has been investigated in Hepatitis C virus (HCV) dynamics (20), but it has not been looked into in SARS-CoV-2 viral dynamic models. Brown *et al*. has shown that the infectious intracellular virus will increase over time post infection, leading to a larger amount of viral replication proteins, which is the main target site of antiviral drugs (19). Since the target sites outnumbered the amount of active drug molecules, there was less drug effect. Therefore, the probability of drug binding would decrease significantly and further undermine the antiviral efficacy (19), reflected in the increasing EC_50_. The increasing EC_50_ suggested the antiviral efficacy was time-dependent and was less potent when the treatment started late. The underlying reason for the reduced antiviral effect is caused by the increment of viral replication related proteins, the target site of most SARS-CoV-2 antivirals, which is proportional with the intracellular viral burden.

Although the hypothesis that intracellular viral load burden is the driving force of EC_50_ shift remains to be further validated, this study took the first step and provided the insight of the relationship between antiviral activity and treatment initiation time. More attention should be paid in clinical practice to those patients who present to hospital late. Accounting for the reduced drug efficacy when the treatment is initiated at late state of infection may be helpful to optimize the dosage and treatment duration in these patients.

## Conclusion

In this study, we conducted a two-way hybrid verification to prove that late treatment initiation reduces antiviral drug efficacy due to a time-dependent increase in EC_50_. Clinically, our results underscores the importance of adjusting dose level and treatment duration to compensate for the reduced drug efficacy when patients presenting late. Future studies need to be done to further validate if the intracellular level of infection is the drive underlying the EC_50_ shift.

## Materials and methods

### Experimental studies

Antiviral assays with static drug concentrations were performed in 6-well plates seeded with 2 × 10^6^ angiotensin-converting enzyme 2 (ACE2)-A549 cells (21), a generous gift from Shinji Makino at the University of Texas Medical Branch, as previously described(19). Briefly, cells were inoculated with the USA-WA1/2020 SARS-CoV-2 strain (BEI Resources, NR-52281) at a multiplicity of infection (MOI) of 0.03 for one hour. After the incubation period, cell monolayers were washed twice to remove any unbound virus and replaced with 3 mL tissue culture medium. Five drug concentrations were evaluated for both nirmatrelvir and GS-441524, alongside a no-treatment control. Viral supernatants were collected daily for 4 days, clarified by high-speed centrifugation, and frozen at -80°C until the end of the study. Plaque assay was performed on viral supernatants using Vero E6 cells to determine the infectious burden in plaque-forming unit (PFU)/mL. The lower limit of quantification (LLOQ) is 100 PFU/mL. Assays were conducted in duplicate and performed twice on two separate days.

The viral load increase was defined as the difference of viral load between day 0 and day 5. The viral replication inhibition was derived from the relative viral load reduction, by calculating the difference of the viral load increase between treatment group and no-treatment control. The experiment-based EC_50_ was then fitted based on the following equation:

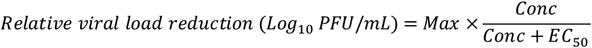

Where *Max* means the maximum relative viral load reduction among all the concentration groups, and *Conc* is the drug concentration.

### Viral dynamic modeling

We considered two candidate models to study the SARS-CoV-2 viral dynamics with varying times of treatment initiation: a TCL model with constant EC_50_ and a TCL model with multiple EC_50_. The two models were then fitted and evaluated with the nirmatrelvir and GS-441524 experimental data.

Both TCL models contain three populations of which the dynamics can be described by four rate constants: susceptible target cells (*T*) get infected and become infected cells (*I*) at a viral infection rate *β, I* then start to produce extracellular free virus (*V*) at a production rate *ρ*, and die at a death rate *δ* after exhausting all the resources. The viral load decline potentially caused by loss of viral infectivity was explained by a rate constant viral clearance *c*. Parameters *δ* and *c* were fixed to 2.58 day^-1^ and 10 day^-1^ according to a published study (7) to deal with the identifiability issue. The TCL model can be described by an ordinary differential equations (ODE) system as below:

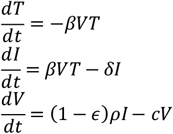

Assuming the viral load at day 0 (V_0_) as initial viral load *V(0)*, the other initial condition was set up according to the experiment design and plausible hypothesis: at *t* = 0, *T*(0) = 2 × 10^6^ and *I*(0) = 0.

Then, for either of these models, a sigmoid maximum possible effect of the drug (E_max_) model was implemented to characterize concentration-effect relationships for nirmatrelvir and GS-441524 data respectively. Given the mechanism of action (MoA) of these two drugs, the antiviral effect (*ε*) on viral production inhibition was applied to viral production rate *ρ*, with *Conc* being the static drug concentration.

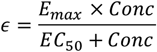

Notably, four EC_50_ values were estimated in the TCL model with multiple EC_50_. Each treatment initiation time had its own EC_50_, instead of sharing one EC_50_ for the entire dataset.

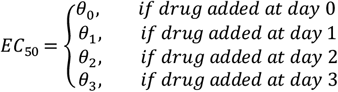

### Software

The model fitting and evaluation were performed using *nlmixr2* package (version 4.0.0) and *rxode2* package (version 4.0.2) in RStudio (R version 4.5.1). Source code was compiled using GCC (GNU Compiler Collection) version 14.2.0. The estimation was performed using the first-order conditional estimation with interaction (FOCEi) method. Censored data which was below the LLOQ was dealt with using Beal’s M3 method (22). The final model was selected based on Bayesian information criterion (BIC), goodness of fit (GOF) plots, and individual plots.

### Simulation of viral dynamics with different antiviral therapies

To investigate the effect of different antiviral therapies, nirmatrelvir – the first-line treatment for SARS-CoV-2 infection – was used as an example drug. A published nirmatrelvir/ritonavir population PK model (18) was used to generate dynamic nirmatrelvir concentration profiles. The simulated PK data was then linked to the developed time-dependent TCL model to simulate the viral load profile under different scenarios. In each scenario, the dose and treatment initiation time were varied to investigate their impact on antiviral effectiveness by comparing the simulated viral load trajectories.

