## Supplementary materials for "Late treatment initiation leads to reduced antiviral potency"

**Supplementary material**

**Supplement 1: Nirmatrelvir PK simulation**

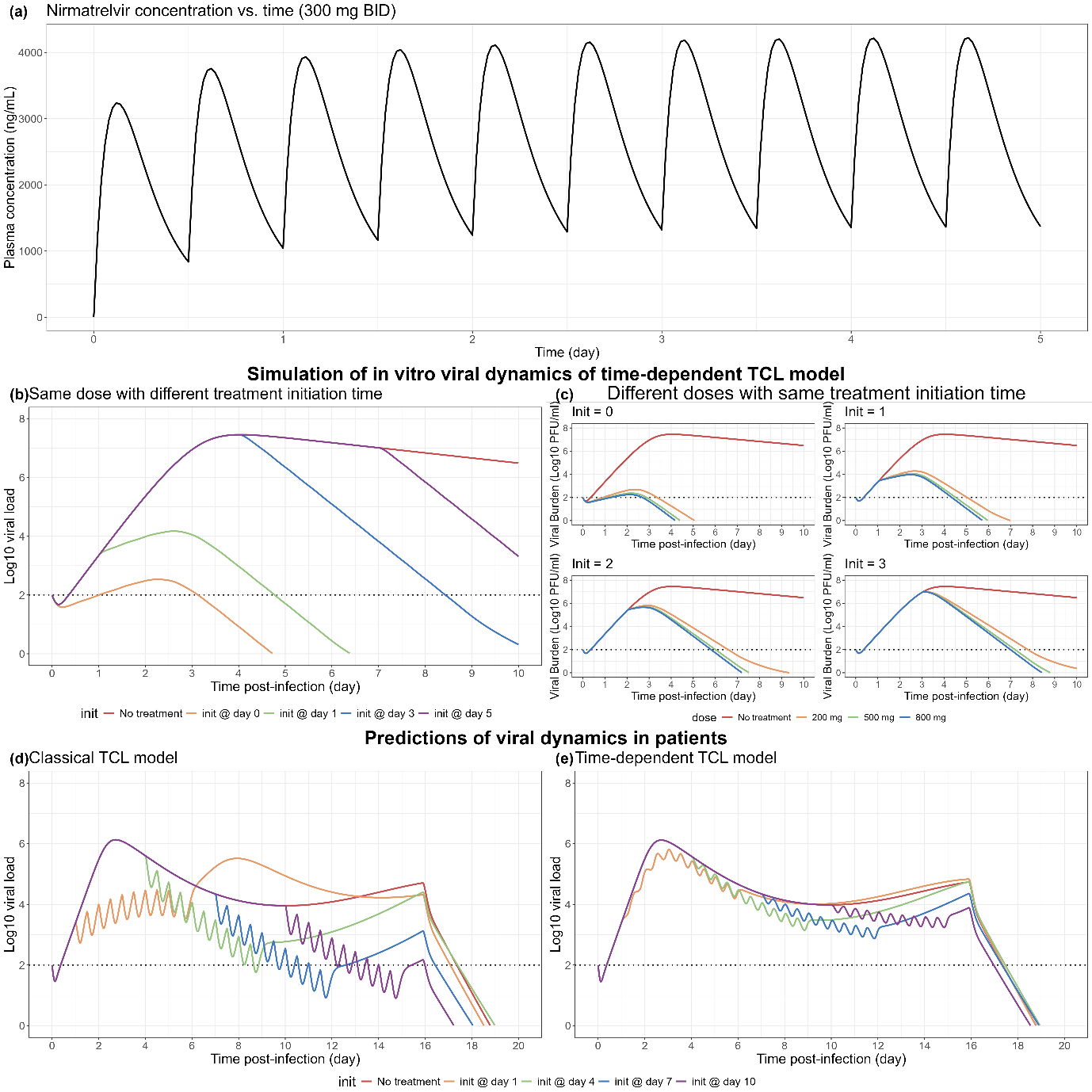

**Fig. S1** Plasma concentration profile given by recommended nirmatrelvir dosing regimen: 300 mg oral dose, twice per day for 5 days.

**Supplement 2: Model code**

```
########################## Environment set-up ############################
####--------------------- Initialization -----------------------####
rm(list=ls())

####--------------------- Import library -----------------------####
library(nlmixr2)
library(dplyr)
library(ggplot2)
library(deSolve)
library(tidyverse)
library(ggpubr)
library(lattice)
library(xpose.nlmixr2)
library(magrittr)
library(tidyvpc)
library(rxode2parse)

####--------------------- Define functions ---------------------####
create_NM_dataset <- function(drug_name, conc_list, trial_no){
  sub_df <- df %>% subset(Drug == drug_name) # Subset df based on drug

  # Convert data sets into NM format
  sub_df <- sub_df %>%
    mutate(EVID = 0, AMT = 0) %>%
    rename(TIME = Time,
           DOSE = Conc,
           DV = Mean,
           INIT = Init,
           GRP = Trial) %>%
    select(TIME,EVID,AMT,DV,DOSE,INIT,GRP)

  # Create an event table
  init_list <- c(0,1,2,3)
  evTable <- data.frame(matrix(ncol = 7,nrow = 0, dimnames = list(NULL, c('TIME', 'EVID', 'AMT', 'DV', 'DOSE', 'INIT', 'GRP'))))

  for (trial in 1:trial_no){
    for (conc in conc_list) {
      for (init in init_list){
        evTable <- evTable %>% add_row(TIME = init,
                                       EVID = 101,
                                       AMT = conc,
                                       DV = 0,
                                       DOSE = conc,
                                       INIT = init,
                                       GRP = trial)
      }
    }
  }

  # Combine two tables and add ID column
  sub_df <- rbind(evTable, sub_df)
  ID_no <- 24*trial_no
  sub_df <- sub_df %>%
    arrange(GRP,INIT,DOSE,TIME) %>%
    mutate(ID = rep(1:ID_no, each = 7)) %>%
    mutate(DOSE = as.factor(DOSE)) %>%
    select(ID,TIME,EVID,AMT,DV,DOSE,INIT,GRP)

  # Add V0
  for (i in 1:ID_no) {
    log_v0 <- sub_df[sub_df$ID == i & sub_df$TIME == 0 & sub_df$EVID == 0, 5]
    sub_df$v0[sub_df$ID == i] <- 3*(10^log_v0)
  }

  return(sub_df)
}
########################## Data pre-processing ############################
# Import dataset
df <- read.csv('data/SARS_CoV_2_tidy.csv',header = TRUE, sep = ',', na.strings = '.')

# Calculate mean viral load
df <- df %>%
  rowwise() %>%
  mutate(Mean = ifelse(is.na(Rep1) & !is.na(Rep2), Rep2,
                       ifelse(!is.na(Rep1) & is.na(Rep2), Rep1, log10((10^Rep1 + 10^Rep2)/2)))) %>%
  mutate(Mean = round(Mean, 3))

# Initialization
conc_NMV <- c(0,0.004,0.0156,0.0625,0.25,1)
conc_RDV <- c(0,2,3,4,6,8)

# Split data set by drug name

NM_df_NMV <- create_NM_dataset(drug_name = 'Nirmatrelvir', conc_list = conc_NMV, trial_no = 2)
NM_df_NMV_CENS <- NM_df_NMV %>% mutate(CENS = ifelse(DV == 1.699, 1, 0),
                                       DRUG = 0)

NM_df_RDV <- create_NM_dataset(drug_name = 'Remdesivir', conc_list = conc_RDV, trial_no = 2)
NM_df_RDV_CENS <- NM_df_RDV %>% mutate(CENS = ifelse(DV == 1.699, 1, 0),
                                       DRUG = 1)

NM_df_all <- bind_rows(NM_df_NMV_CENS, NM_df_RDV_CENS) %>%
  mutate(ID = rep(1:96, each = 7))

############################### Define the model ##############################
####------------------- TCL + single EC50 ------------------####
TCL_single_EC50 <- function(){
  # Set initial values
  ini({
    # infection rate
    tbeta <- log(1*10^-8)
    # death rate of infected cells
    tdelta <- fix(log(2.58)) # delta (1/day)
    # viral production rate
    trho <- log(1000) # rho (PFU/cell/day)
    # viral clearance
    tc <- fix(log(10)) # c (1/day)
    # EC50
    tec50_NMV <- log(c(0,0.1))
    tec50_RDV <- log(c(0,2))
    temax <- fix(0)

    # residual error
    add.err <- 1
  })
  model({
    # individual values
    beta <- exp(tbeta)
    delta <- exp(tdelta)
    rho <- exp(trho)
    c <- exp(tc)
    emax <- exp(temax)
    ec50_NMV   <- exp(tec50_NMV)
    ec50_RDV   <- exp(tec50_RDV)

    # initial state
    T(0) = 2*10^6
    I(0) = 0
    V(0) = v0

    # ODE
    # PK
    d/dt(center) = 0
    Cp = center

    # Single EC50
    if (DRUG == 0) {
      ec50 <- ec50_NMV
    } else {
      ec50 <- ec50_RDV
    }

    E = (emax*Cp)/(ec50+Cp)

    d/dt(T) <- -beta*V*T
    d/dt(I) <- beta*V*T - delta*I
    d/dt(V) <- (1-E)*rho*I - c*V

    VL = log10(V/3)
    VL ~ add(add.err)
  })
}
####------------------- TCL + multi EC50 ------------------####
TCL_multi_EC50 <- function(){
  # Set initial values
  ini({
    # infection rate
    tbeta <- log(1*10^-8)
    # death rate of infected cells
    tdelta <- fix(log(2.58)) # delta (1/day)
    # viral production rate
    trho <- log(1000) # rho (PFU/cell/day)
    # viral clearance
    tc <- fix(log(10)) # c (1/day)
    # EC50
    tec50_NMV_0 <- log(c(0,0.1))
    tec50_NMV_1 <- log(c(0,0.1))
    tec50_NMV_2 <- log(c(0,0.1))
tec50_NMV_3 <- log(c(0,0.1))

    tec50_RDV_0 <- log(c(0,2))
    tec50_RDV_1 <- log(c(0,2))
    tec50_RDV_2 <- log(c(0,2))
tec50_RDV_3 <- log(c(0,2))

    temax <- fix(0)

    # residual error
    add.err <- 1
  })
  model({

    # individual values
    beta <- exp(tbeta)
    delta <- exp(tdelta)
    rho <- exp(trho)
c <- exp(tc)
    emax <- exp(temax)
    ec50_NMV_0 <- exp(tec50_NMV_0)
    ec50_NMV_1 <- exp(tec50_NMV_1)
    ec50_NMV_2 <- exp(tec50_NMV_2)
    ec50_NMV_3 <- exp(tec50_NMV_3)
    ec50_RDV_0 <- exp(tec50_RDV_0)
    ec50_RDV_1 <- exp(tec50_RDV_1)
    ec50_RDV_2 <- exp(tec50_RDV_2)
    ec50_RDV_3 <- exp(tec50_RDV_3)

    # initial state
    T(0) = 2*10^6
    I(0) = 0
    V(0) = v0

    # ODE
    # PK
    d/dt(center) = 0
    Cp = center

    # Multi EC50 for both drugs
    if (DRUG == 1 & INIT == 0) {
      ec50 <- ec50_RDV_0
    } else if (DRUG == 1 & INIT == 1){
      ec50 <- ec50_RDV_1
    } else if (DRUG == 1 & INIT == 2){
      ec50 <- ec50_RDV_2
    } else if (DRUG == 1 & INIT == 3){
      ec50 <- ec50_RDV_3
    } else if (DRUG == 0 & INIT == 0){
      ec50 <- ec50_NMV_0
    } else if (DRUG == 0 & INIT == 1){
      ec50 <- ec50_NMV_1
    } else if (DRUG == 0 & INIT == 2){
      ec50 <- ec50_NMV_2
    } else {
      ec50 <- ec50_NMV_3
    }

    E = (emax*Cp)/(ec50+Cp)

    d/dt(T) <- -beta*V*T
    d/dt(I) <- beta*V*T - delta*I
    d/dt(V) <- (1-E)*rho*I - c*V

    VL = log10(V/3)
    VL ~ add(add.err)
  })
}
################################# Model fitting ###################################

run_res_singleEC50 <- nlmixr(TCL_single_EC50, NM_df_2, est = "focei", control = list(print=0), outerOpt="bobyqa")

run_res_multiEC50 <- nlmixr(TCL_multi_EC50, NM_df_2, est = "focei", control = list(print=0), outerOpt="bobyqa")
```
